# Life with only 28 tRNAs: Reduced translation accuracy compensates for the lack of twelve tRNAs in *Salmonella enterica*

**DOI:** 10.1101/2024.04.04.588050

**Authors:** Joakim Näsvall

**Affiliations:** Dept. of Medical Biochemistry and Microbiology, Uppsala University, Uppsala, Sweden

## Abstract

Despite the link between codon usage bias and the composition of the tRNA pool, the evolutionary forces shaping codon usage and tRNA pools remain largely untested by experiment. This study investigates the relationship between tRNA pool composition and synonymous codon usage (SCU) by deleting twelve nonessential tRNAs in *Salmonella enterica*, generating an organism (ΔT12) with 28 essential tRNAs left, and a severe imbalance between its tRNA pool and SCU. Mutations selected during the construction of ΔT12 and in subsequent evolution experiments, suggest two key mechanisms for compensating the fitness effects of this imbalance: (*i*) Near-cognate tRNA adaptation: mutations or gene copy number variations allowed remaining tRNAs to better read codons originally assigned to missing tRNAs. (*ii*) Reduced translation accuracy: mutations in ribosomal proteins (S3, S4, S5, L7/L12) and EF-Tu likely increased the rate of near-cognate decoding, prioritizing translation speed over accuracy. These findings suggest that translation rate may be a stronger evolutionary pressure than maintaining perfect accuracy when there is a mismatch between the tRNA pool and SCU. The ΔT12 strain provides a valuable tool for further exploring the co-evolution of the tRNA pool and SCU.

## Introduction

The genetic code exhibits degeneracy at three levels (Figure 1): (*i*) multiple codons for a single amino acid, (*ii*) some tRNAs reading multiple codons for the same amino acid, and (*iii*) decoding of some codons by multiple tRNAs. The latter degeneracy creates a layer of redundancy, as translation could continue (albeit potentially inefficiently or inaccurately) even without a specific tRNA.

**Figure 1.**
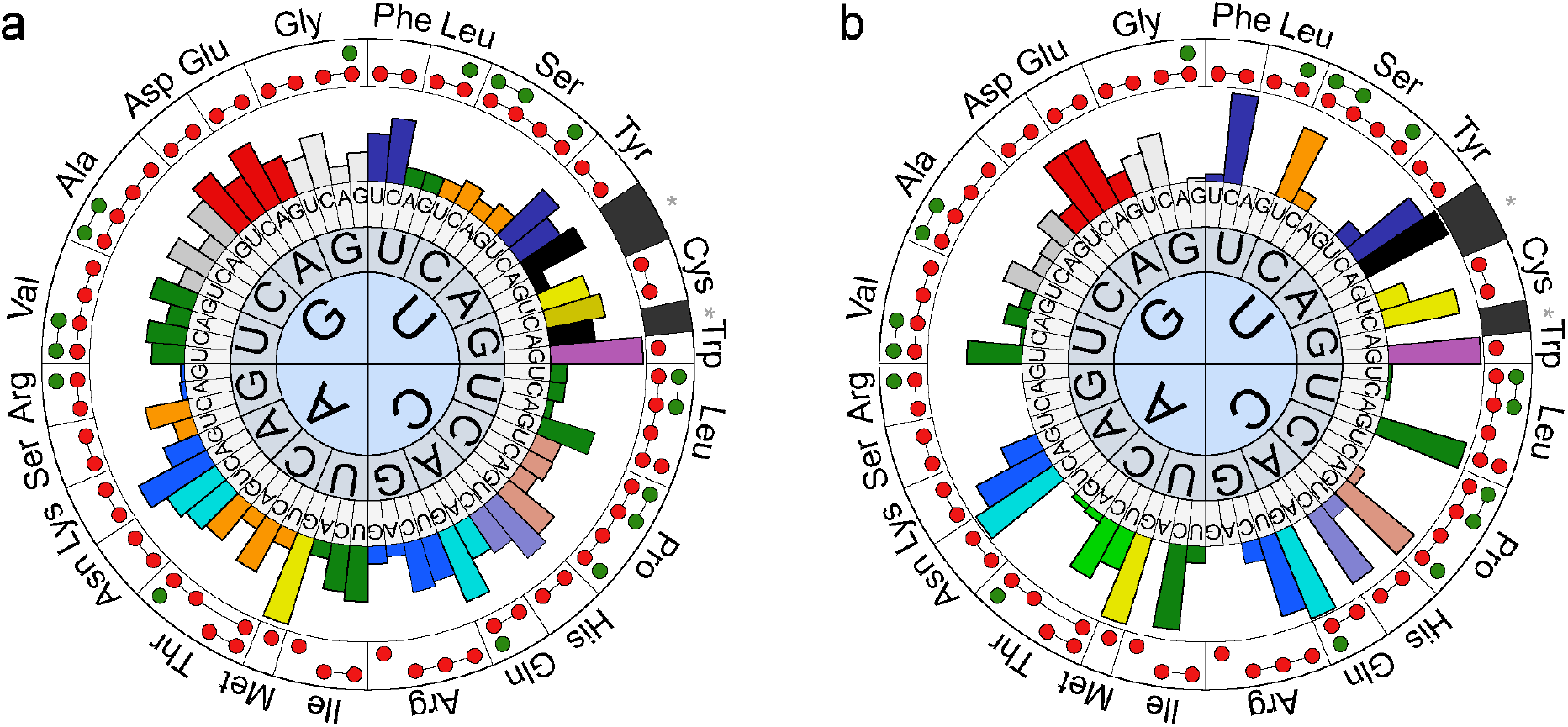
Degeneracies, redundancies, and biases in the genetic code. The three inner circles represent the 64 codons in the genetic code, bars represent relative synonymous codon usage (rSCU). Symbols in the outer circle represent the tRNAs (40 unique anticodons, excluding the initiator tRNA^fMet^ and the selenocysteine tRNA^sec^) encoded in the genome. Two or more dots connected by a line indicate a tRNA able to read several synonymous codons, while single dots indicate tRNAs capable of reading only one codon. Green dots represent the 12 nonessential tRNAs, red dots represent essential tRNAs. **(a)** rSCU in all coding sequences of *Salmonella enterica*. **(b)** rSCU in the two genes encoding EF-Tu (one of the most highly expressed proteins) in *S. enterica*. rSCU is from the Codon Usage Database (https://www.kazusa.or.jp/codon/), tRNA content from GtRNAdb (http://gtrnadb.ucsc.edu/GtRNAdb2/). The decoding specificities of the tRNAs and the rationale for which tRNAs were considered potentially nonessential are explained in Supplementary Text and Figure S1.

The decoding specificities of tRNAs are primarily determined by anticodon sequences, but other factors like nucleotide modifications at the wobble position further refine it^1^. For example, tRNAs with the modified nucleoside uridine-5-oxyacetate (cmo^5^U) at the wobble position of the anticodon can read codons ending with A, G, U, and sometimes C^2–4^, while tRNA^Ile^_k2C_ with lysidine (k^2^C; a modified cytidine) can read only the AUA codon^5^.

Synonymous codon usage (SCU) varies between organisms, and correlates with maximum growth rate, the genome’s GC content, and the composition of the tRNA pool^6^. Notably, SCU also varies within a single genome. Highly expressed genes exhibit a strong bias towards using specific codons and avoiding others, while genes with low expression levels tend to have a more random codon usage. A plausible explanation for the evolution of codon usage bias is that selection favors a balance between tRNA availability and SCU. This ensures that the tRNAs corresponding to the most abundant codons are also the most plentiful^7^. Certain codons prone to misreading or frameshifting are disfavored. Additionally, tRNA expression is balanced to not only ensure the translation of their cognate codons but also to minimize misreading of other codons^8^.

Several hypotheses explain the evolution of SCU: mutational biases, selection for translation rate or accuracy, or a combination of both^9^. Additional levels of selection are suggested to add to the balance between SCU and the composition of the tRNA pool, *e*.*g*. there is a tendency for organisms inhabiting the same environment to have similar tRNA pools and SCU, an effect which is suggested to be caused by frequent horizontal gene transfer^10^. Horizontally transferred genes with a SCU that is not adapted to the tRNA pool of the new host organism are costly and poorly expressed. This cost can be ameliorated both in laboratory experiments and in nature by altering the composition of the tRNA pool^11,12^. Thus, it is plausible that an imbalance between codon usage and tRNA availability (*e*.*g*., due to horizontally transferred DNA) are compensated either by altering the codon usage to match the tRNA pool, or to alter the tRNA pool to match the new DNA, *e*.*g*. by tRNA gene copy number variation and evolution of new tRNAs that are better adapted for efficient and/or accurate translation of the new codon usage.

To investigate the selection pressures shaping the tRNA pool, twelve nonessential tRNAs were deleted in *Salmonella enterica*, leaving it with 28 essential tRNAs. This minimal pool resembles the tRNA pools found in organisms with minimal genomes and codon usage biases that are very different from *S. enterica*. As this organism still has the original codon usage of *S. enterica*, its tRNA pool is far from balanced to its SCU. Laboratory evolution revealed two main compensatory pathways: (*i*) mutations and expression changes in remaining tRNAs, potentially enhancing near-cognate decoding efficiency, and (*ii*) mutations in ribosomal proteins known to reduce translation accuracy, thereby increasing near-cognate decoding. These findings provide valuable insights into the intricate relationship between codon usage bias and the composition of the tRNA pool.

## Results and Discussion

### Twelve tRNA species are nonessential in *S. enterica*

Our previous work demonstrated that five tRNA species, present in family codon boxes (groups of codons encoding the same amino acid), were nonessential in *S. enterica*^2,3^. This implied that other tRNAs for the same amino acids could read codons beyond what Crick’s wobble hypothesis predicted^14^. Based on this updated understanding, eight additional potentially nonessential tRNAs, whose cognate codons could potentially be read by another tRNA specifying the same amino acid, were identified (Figure S1; Table 1).

**Table 1.**
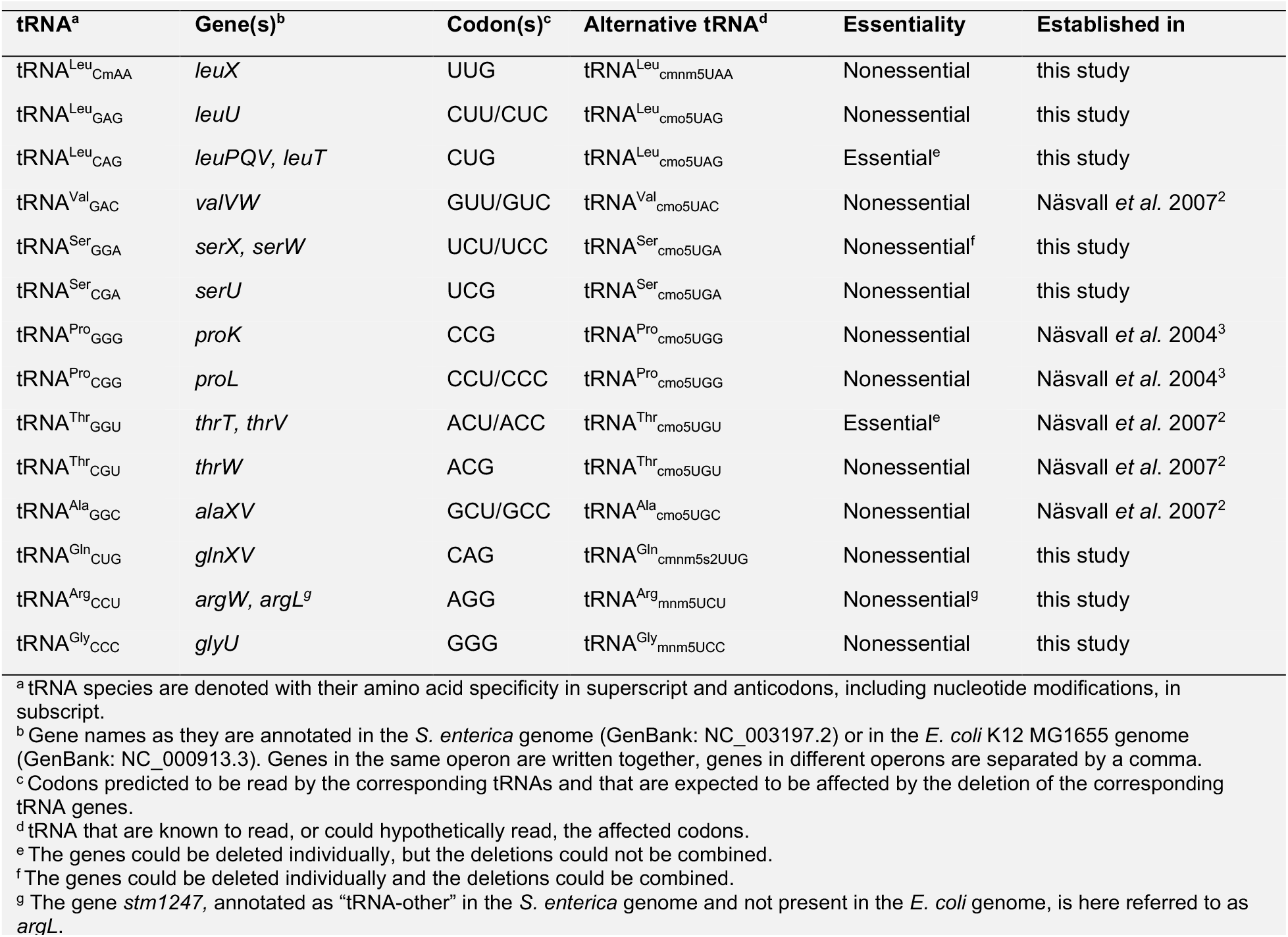
Of fourteen potentially redundant tRNAs, twelve are nonessential in *S. enterica*.

To test these predictions, DIRex, a method enabling single-step, scar-free deletions^15,16^ was used. DIRex facilitates the transfer of mutations between strains via transduction and allows for simple marker recycling, enabling the repeated use of the same selection marker (Figure 2). A total of thirteen tRNAs were targeted for deletion: eight novel candidates predicted to be nonessential and five previously demonstrated as such^2^. Ten of these tRNAs are encoded by single genes or tandem duplicates within a single locus. The remaining three require a combined deletion of two genes or operons located in different loci for complete removal. This resulted in deletions of totally 21 genes across 16 individual loci.

**Figure 2.**
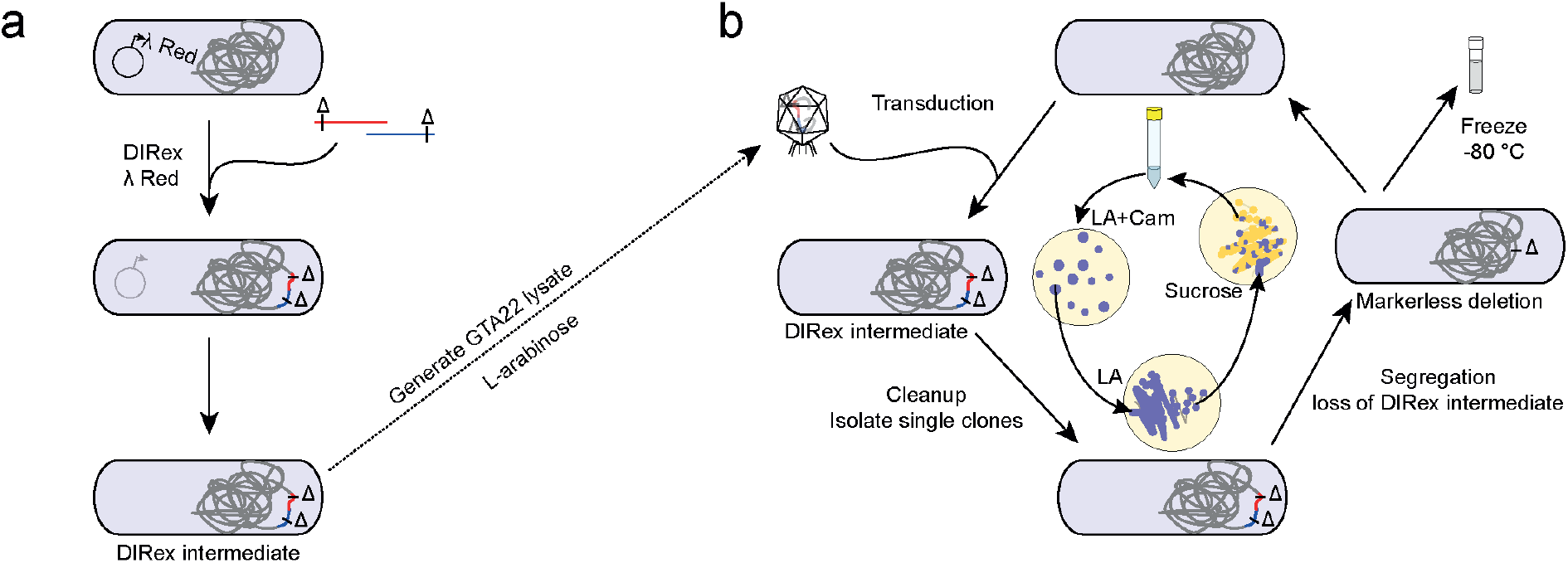
Strategy for deleting multiple genes. **(a)** Individual deletions were constructed with DIRex. Two PCR products, with overlapping parts of a selectable and counter-selectable cassette *(AcatsacA;* conferring chloramphenicol resistance, sucrose sensitivity, and blue color, containing at each end identical but inversely oriented 800 bp sequences) and 5’-extensions that generate a direct repeat containing the deletion junction, were co-transformed into bacteria expressing λ Red. Chloramphenicol resistant transformants contain a DIRex intermediate with directly repeated 30 bp sequences containing the designed deletion junction flanking the *AcatsacA* cassette. **(b)** Transduction cycles to introduce the deletions one after another. Each cycle starts with transduction of a DIRex intermediate into a recipient strain using a lysate generated by inducing the GTA22 prophage in a donor strain. Chloramphenicol resistant, blue (and sucrose sensitive) transductants were streaked for single colonies once on LA plates to generate pure clones. Colonies of pure, blue clones from LA plates were streaked once on sucrose selection plates to select loss of the DIRex intermediate and generation of the final deletion (sucrose resistant, colorless, chloramphenicol sensitive) before being used as recipient in the next transduction cycle.

One tRNA, tRNA^Leu^_CAG_, was identified as essential. Although both loci (four genes) encoding this tRNA were targeted for deletion, and individual deletions in each locus were viable, a strain containing deletions in both loci could not be obtained. Consequently, twelve out of the fourteen predicted nonessential tRNAs are indeed nonessential in *S. enterica*. The essentiality of tRNA^Leu^_CAG_ is not unexpected, considering it recognizes the most abundant leucine codon (>50% of all leucine codons) and is itself one of the most abundant tRNAs^17^.

While deletion of twelve tRNAs confirmed their nonessentiality, three deletions (*alaWX, valVW*, and *leuU*) resulted in slow growth and temperature sensitivity (Figure 3a). This suggests that although these tRNAs are dispensable for viability, their absence impairs optimal growth.

**Figure 3.**
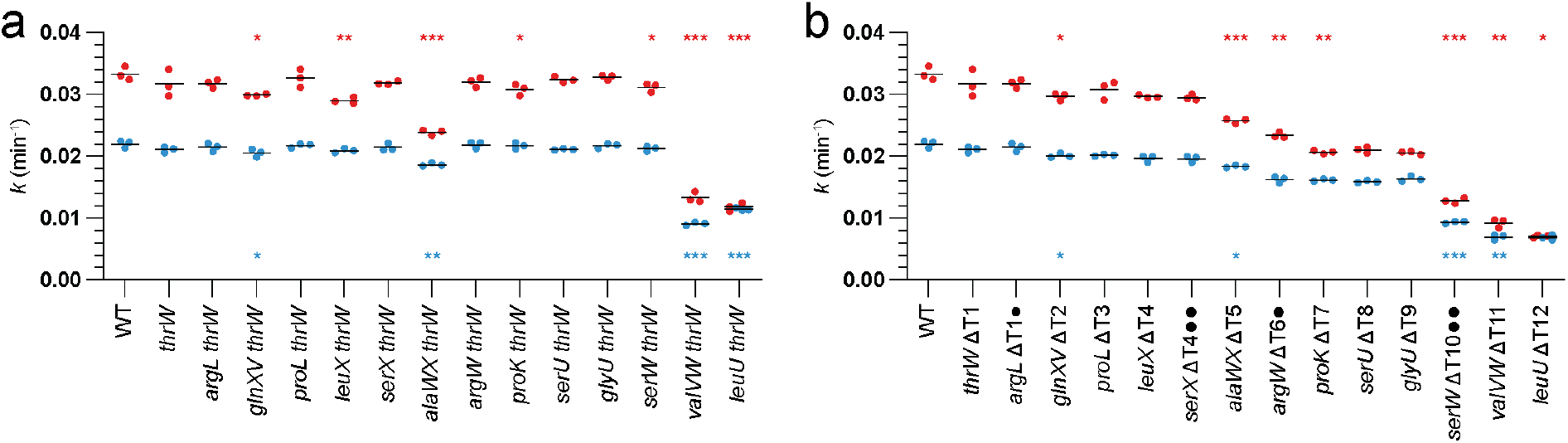
Growth rates of tRNA deletion mutants at 37 °C (red) and 30 °C (blue). **(a)** Individual deletions. The deletions are all combined with Δ*MhrW* (that was deleted as part of constructing GTA22 which is present in all strains except the WT). **(b)** The construction series leading up to ΔT12. The mutants are denoted by the most recently deleted tRNA gene followed by an indication of the number of missing tRNAs; *e*.*g*., ΔT1 is missing one tRNA, ΔT2 is missing two, *etc*.*•, argL* and *argW* encode tRNAs with the same anticodons; ••, *serX* and *serW* encode identical tRNAs. Only completely missing tRNAs (all genes encoding tRNAs with the same anticodon deleted) add to the count of missing tRNAs. The three leftmost strains in (a) and (b) are the same. Asterisks show significant differences according to a two-tailed Student’s *t*-test; (a) compared to the *thrW* deletion strain, (b) each mutant compared to its parent *(i*.*e*., ΔT12 was compared with ΔT11, which in turn was compared to ΔT10, *etc*.), ^*^, *p* < 0.05; ^**^, *p* < 0.005; ^***^, *p* < 0.005.

To combine all twelve deletions of nonessential tRNAs into a single viable strain, a sequential generalized transduction approach was employed, introducing the deletions one after another (Figure 2b; Figure S2). To expedite each transduction cycle and avoid selection for phage-resistant mutants, strains harboring the arabinose-inducible artificial gene transfer agent GTA22^18^ were utilized both as donors and recipients. This prophage is constructed from a P22 prophage by deletions at both the prophage ends, also deleting the attachment site in the *thrW* (tRNA^Thr^_CGU_ ) gene, rendering strains containing GTA22 already deleted for *thrW*.

Following the 13th and final transduction cycle, single clones from 9 different lineages that had been kept separate for 2 – 4 transduction cycles were selected for whole-genome sequencing (WGS; Figure S2). While all sequenced clones harbored the intended tRNA deletions, 6-7 unintended mutations were also identified in all lineages (Table S1). Of these mutations, six were shared between all sequenced strains, including three mutations potentially affecting translation (*dksA*[ΔN88-R89], *leuP*[tRNA^Leu^_CAG_; G18T], and a copy number variation within the *serV*-*argVYZQ* tRNA operon that results in one fewer gene encoding tRNA^Arg^_ICG_; Δ*argZQ*-DUP*argY*). WGS analysis of clones from transduction cycle 10 revealed that the six common mutations were already present in all lineages at that point. One of the clones (DA67309; hereafter ΔT12) containing only the six common mutations was selected for further experiments. In a separate series of transductions, another mutant lacking the same twelve tRNAs was generated (hereafter ΔT12b; Figure S3). WGS of this strain revealed a mutation affecting one of the two genes encoding EF-Tu (*tufA* [I367Leu]; Table S2).

The growth rates of strains harboring combinations of deletions (Figure 3b) exhibited a generally declining trend with each successive deletion, and the final ΔT12 strain was unable to grow on plates at 37 °C. However, some intermediate strains, particularly following the introduction of the *alaWX* deletion at cycle 6, displayed a less pronounced growth reduction compared to strains with individual deletions. This observation suggests that some unintended mutations may have been selected during the construction process, acting as compensatory mutations. The *leuP* mutation, if it affects the accuracy of the encoded variant of tRNA^Leu^_CAG_, could potentially compensate for the loss of *leuX* (encoding tRNA^Leu^_CmAA_ ) by enabling the misreading of its near-cognate codon UUG through a first-position mismatch. DksA is a co-regulator of the stringent response^19^. Whatever the specific effects of the observed mutation (overactivity, complete loss, or partial loss of function) it is likely to influence the transcription of ribosomal components and tRNAs, but the effects of this mutation are difficult to predict. In the separate transduction series, the final ΔT12b strain was viable at 37 °C, possibly due to the EF-Tu-mutation which was selected after the introduction of one of the more severe deletions (Figure S3).

To elucidate the mechanisms underlying the severe fitness reduction observed in ΔT12, evolution experiments were conducted by serial passage in batch cultures. Sixteen populations were propagated at 30°C and another sixteen at 37°C for 245 generations. Notably, all populations displayed improved growth compared to ΔT12 at the endpoint, and the prior growth temperature resulted in no obvious difference (Figure S4).

Whole-genome sequencing (WGS) of twenty populations was used to identify the mutations responsible for compensating for the growth defects in ΔT12 (Table 2). This analysis revealed 1 – 2 additional mutations in each population, with a striking observation: all populations harbored mutations in genes associated with translation (Table 2). These mutations involved ribosomal small subunit proteins S3 (*rpsC*; 1 population), S4 (*rpsD*; 11 populations), S5 (*rpsE*; 7 populations), large subunit protein L7/L12 (*rplL*; 2 populations), and duplications of the *serT* gene (encoding tRNA^Ser^_cmo5UGA_; 2 populations).

**Table 2.** Candidate compensatory mutations in evolved populations. See Table S3 for more details.

| Strain | Population<sup>a</sup> | WGS<sup>b</sup> | Candidate mutation(s)<sup>c</sup> | Linkage<sup>d</sup> |  |
| --- | --- | --- | --- | --- | --- |
|  |  |  |  | *rplQ* | *rplK* |
| DA67309 | ΔT12 | Yes | - | - | - |
| DA75235 | 30:1 | Yes | *rpsE* (R112S) | Yes | Nd |
| DA75236 | 30:2 | Yes | *rpsD* (Y204fs) | Yes | Nd |
| DA75237 | 30:3 | Yes | *rpsD* (Y204fs) | Yes | Nd |
| DA75238 | 30:4 | Yes | *rpsE* (G109D), dup(*yccW-hpaF*; *serT*) | Yes | Nd |
| DA75239 | 30:5 | Yes | *rpsD* (L191fs), dup(*pepN-hpaI*; *serT*) | Yes | Nd |
| DA75240 | 30:6 | Yes | *rpsE* (A99V) | Yes | Nd |
| DA75241 | 30:7 | Yes | *rpsE* (V123F) | Yes | Nd |
| DA75242 | 30:8 | Yes | *rpsD* (del13bp; L199-L203_ fs) | Yes | Nd |
| DA75243 | 30:9 | Yes | *rpsE* (A99D) | Yes | Nd |
| DA75244 | 30:10 | Yes | *rpsC* (R127H), *rplL* (del V41-A42) | Yes | Yes |
| DA75245 | 30:11 | No | - | Yes | Nd |
| DA75246 | 30:12 | Yes | *rpsE* (A139E), *rplL* (del V41-A42) | No<sup>e</sup> | No<sup>e</sup> |
| DA75247 | 30:13 | Yes | *rplL* (del G44-A49) | No | Yes |
| DA75248 | 30:14 | Yes | *rpsD* (del8bp; N196fs) | Yes | Nd |
| DA75249 | 30:15 | Yes | *rpsD* (del13bp; L199-L203_ fs) | Yes | Nd |
| DA75250 | 30:16 | No | - | Yes | Nd |
| DA75251 | 37:1 | Yes | *rpsD* (D194fs) | Yes | Nd |
| DA75252 | 37:2 | Yes | *rpsD* (*207Q; +37aa) | Yes | Nd |
| DA75253 | 37:3 | No | - | Yes | Nd |
| DA75254 | 37:4 | No | - | Yes | Nd |
| DA75255 | 37:5 | Yes | *rpsE* (T103P) | Yes | Nd |
| DA75256 | 37:6 | No | - | Yes | Nd |
| DA75257 | 37:7 | Yes | *rpsD* (K206T) | Yes | Nd |
| DA75258 | 37:8 | No | - | Yes | Nd |
| DA75259 | 37:9 | Yes | *rpsD* (D190fs) | Yes | Nd |
| DA75260 | 37:10 | No | - | Yes | Nd |
| DA75261 | 37:11 | No | - | Yes | Nd |
| DA75262 | 37:12 | Yes | *rpsD* (S205F) | Yes | Nd |
| DA75263 | 37:13 | No | - | Yes | Nd |
| DA75264 | 37:14 | No | - | Yes | Nd |
| DA75265 | 37:15 | No | - | Yes | Nd |
| DA75266 | 37:16 | No | - | Yes | Nd |

<sup>a</sup>Populations are named according to what temperature they were evolved at (30 or 37 °C) and are numbered 1 – 16. For growth rates of the evolved populations, see Figure S4.
<sup>b</sup>"Yes", the strain was whole genome sequenced; "No" it was not.
<sup>c</sup>Candidate mutations identified from WGS.
<sup>d</sup>Linkage of compensating mutation to *rplQ* or *rplK* as determined by transductions. Linkage was determined by the presence of two types of transductants with distinguishable colony sizes. Nd, not determined.
<sup>e</sup>No obvious difference in colony sizes among the transductants despite expected linkage.

The most likely candidate for reading codons UCU, UCC, and UCG in the absence of tRNA^Ser^_GGA_ and tRNA^Ser^_CGA_ is the tRNA encoded by *serT*. Overexpression of *serT* is expected to compensate for the lack of the other serine tRNAs. Conversely, mutations in ribosomal proteins S4 (including many C-terminal truncations), S5, and L7/L12^20–24^ (including in-frame deletions similar to those identified in this study) are known to decrease translational accuracy.

Transductions with selection markers positioned near the genes of interest were used to directly connect the identified mutations to the fitness compensation observed in ΔT12. This approach also allowed testing for the presence of compensatory mutations in the same genes in the non-sequenced populations. Phage P22 (and consequently GTA22) can co-transduce nearby genes along with the selected marker, at a frequency that decreases with increasing distance from the marker. Two duplication-insertions (Dup-Ins)^16,25^ were generated – one surrounding the *rplQ* gene (linked to *rpsD, rpsE*, and *rpsC* at distances of 7.5 kb, 3.8 kb, and 1.5 kb, respectively) and another surrounding *rplK* (adjacent to *rplL* at 2.5 kb).

Following transduction, a fraction of the transductants were expected to inherit the wild-type allele from the donor, reverting them to slow growth. The remaining fraction would retain the mutant allele from the recipient, thereby maintaining the faster growth phenotype. In all but one case (Table 2), these transductions confirmed the involvement of candidate mutations in the ribosomal protein genes as contributing factors to the observed compensation. The single population where linkage could not be determined harbored mutations linked to both markers and several other low frequency mutations affecting translation, which could have masked the effects of reverting one of the mutations. Furthermore, the results from the non-sequenced populations suggest that all harbor compensatory mutations linked to *rplQ*, implying *rpsD, rpsE*, or *rpsC* as likely targets.

The ΔT12b strain was also evolved (Table S4) at 37 °C for 300 generations. Sequencing of eight evolved populations revealed mutations in *rpsD* (2 populations), *rpsE* (6 populations), *leuQ* (1 population), and *leuT* (2 populations). In order to gain more insight, a mutator derivative of ΔT12b was also evolved and eight populations were sequenced (Table S5). This resulted in mutations in *rpsE* (2 populations), *rpsC* (3 populations), and *leuW* (4 populations). The tRNA genes *leuQ* and *leuT* both encode tRNA^Leu^_CAG_, and *leuW* encode tRNA^Leu^_cmo5UAG_ . The mutations altered the anticodon to GAG (*leuQ, leuT*, one population each) and AAG (1 *leuT* mutation and all four *leuW* mutations). Thus, the tRNA^Leu^ mutations are expected to result in improved reading of the CUC and CUU codons, compensating for the missing tRNA^Leu^_GAG_ (*leuU*).

## Conclusions

The viability of a bacterial strain lacking twelve out of fourteen “redundant” tRNAs presents a valuable tool for investigating the evolutionary forces that shape the balance between tRNA pool composition and codon usage bias. The compensatory mutations observed during the construction and subsequent evolution of ΔT12 suggest two classes of compensatory mechanisms. One class specifically compensates for the loss of individual tRNAs, by altering the sequence or gene copy number (and thus expression) of tRNAs that are near-cognate to the missing tRNAs. The other class consists of mutations that affect translation accuracy, potentially facilitating faster near-cognate decoding of “starving” codons. Interestingly, these findings suggest that translation accuracy due to misincorporation at “starving” codons may not be the primary concern when cognate tRNAs are missing or present at suboptimal levels. Instead, the results favor translation rate as a more significant selection pressure in the co-evolution of tRNA content and codon usage.

## Materials and Methods

### Strains and media

All bacterial strains are listed in Table S6. As liquid medium, Lysogeny broth (LB; 10 g/L Tryptone [Sigma], 5 g/L yeast extract [Sigma], 10 g/L NaCl) was used. To make LB agar (LA) plates, LB was supplemented with 15 g/L Bacto agar (Sigma). For transferring chromosomal markers between strains, generalized transduction with phage P22 HT 105/1 *int-201*26 or GTA22^18^. Briefly, GTA22 was constructed by introducing a copy of the P22 antirepressor under the arabinose inducible *ParaBAD* promoter and deleting genes from both ends of a P22 *sieA44* HT 105/1 prophage. Notably, the deletions remove the *thrW* tRNA gene that serves as the attachment site for P22, the integrase/excisionase genes and the O-antigen modification locus *gtr*. The modifications makes the phage non-virulent and arabinose inducible, while still allowing the host to serve as both donor and recipient in transductions. Deletions were constructed using DIRex as described elsewhere^15,16,18^, using the primers listed in Table S7. Chloramphenicol (12.5 mg/L) was used for positive selection of *Acatsac*-cassettes in DIRex recombineering^15,16^, and sucrose selection medium (salt-free LA with 5 % [w/v] sucrose [Sigma]) was used for counter-selection against *Acatsac* cassettes. As some mutants failed to grow on salt-free sucrose selection plates, 0.5 % (w/v) NaCl was added for some counter-selections to work.

### Construction of ΔT12

To construct a strain lacking the twelve nonessential tRNAs, sequential GTA22 transductions (Figure 2) were used to introduce the deletions one after another in the order indicated in Figure S2. Each transduction cycle consisted of first introducing a tRNA gene deletion with a DIRex intermediate (*AcatsacA*) containing the genes *cat* (chloramphenicol resistance) and *sacB* (sucrose sensitivity) flanked by two identical but inversely oriented *amilCP* genes (blue chromoprotein), selecting chloramphenicol resistance. Transductants were purified once non-selectively before being streaked on sucrose selection plates to select cells that lost the DIRex intermediate. Sucrose resistant, blue colonies were saved and used as recipients in the next transduction cycle. As some of the individual deletions displayed a relatively more severe growth phenotype at 37 °C than at 30 °C (Figure 3), all steps were done at 30 °C. the sucrose selection plates were modified by adding 0.5 g/L NaCl for the last two transduction cycles as the introduction of the Δ*valVW* deletion the transductants were unable to grow on salt-free LA. For the last five transduction cycles the order of the deletions was varied by introducing Δ*serU*, Δ*glyU*, Δ*serW*, Δ*valVW*, and Δ*leuU* in different orders in different lineages, resulting in twelve “final” clones. Nine of these were whole genome sequenced (Table S1). One of the “final” clones (DA67309) was chosen for further experiments.

### Construction of ΔT12b and identification of a potentially compensating *tufA* mutation

To construct another strain lacking the twelve nonessential tRNAs, a similar transduction series as described above, but using P22, was used to introduce the deletions one after another in the order the strains are listed in Figure S3. The starting strain contained a deletion of the *gal* operon promoter Δ(*P*_*gal*_) and a deletion of the *galE* gene (Δ*galE*::*T*_*lux*_), which renders it conditionally P22 resistant (resistant in the absence of galactose, sensitive in the presence). These mutations were included to make cleanup after transductions faster by minimizing the risk of P22 infection in the transductants (but later observations showed that cells grown in LB can still be infected by P22, presumably due to the presence of a low concentration of galactose in LB). For the last three transduction cycles the order of the deletions was varied by introducing Δ*leuU*, Δ*leuX*, and Δ*valVW* in different orders, resulting in six “final” clones that were whole genome sequenced (Table S2). The mutation *tufA* (I367L) was found in all sequenced end-point strains, and through local sanger sequencing of clones from earlier points in the series, the time of acquiring the *tufA* mutation was determined to be at the introduction of the eleventh (Δ*glyU*) deletion, coinciding with an increase in growth rate compared to the parent from the previous transduction cycle (Figure S3). One of the “final” clones (DA59802) was chosen for further experiments.

### Evolution experiment with ΔT12

A total of 32 populations were started by inoculating single colonies of the ancestral strain (DA67309; ΔT12) into 1 mL of LB in 10 mL plastic tubes. All cultures were grown at 30 °C overnight. After this initial cycle at 30 °C, 16 cultures were passaged at 30 °C and the remaining 16 at 37 °C, by 1:5000 dilutions into 5 ml fresh LB every 24 h for 20 cycles (totalling approximately 245 generations).

### Evolution experiments with ΔT12b

Eight derivatives of ΔT12b with a *mutS* mutation (*mutS*(dup500bp)::*Acatsac1*^1^) and eight without the *mutS* mutation (but with GTA22 and the *gal* operon restored) were passaged by 1:5000 dilutions into 5 ml LB every day for 24 cycles (approximately 300 generations), and were then whole genome sequenced (Tables S4 and S5).

### Growth rate determinations

Overnight cultures (in 1 mL LB) were grown from single colonies (constructed strains) or from 1 μL samples of thawed freezer stocks (evolved populations) at 30 °C. Pure clones were grown as three biological replicates, while populations were grown as a single sample from each population. The overnight cultures were diluted 1:1000 into fresh LB, after which 300 μL was transferred into wells on two Bioscreen honeycomb plates. The two plates were incubated in separate Bioscreen C readers, one at 30 °C and one at 37 °C, with continuous moderate shaking and with reading of the optical density at 600 nm (OD_600_) every four minutes for 24 h. For growth rate determinations, the OD_600_ data was plotted against time to find the exponential phase (which ended at OD600 = 0.24 for most strains except DA67309 [ΔT12] for which it ended at OD600 = 0.16 at 30 °C and OD600 = 0.1 at 37 °C, and DA65699 [Δ*leuU* Δ*thrW*] for which it ended at OD600 = 0.15 at 37 °C). The data from the exponential phase was fitted (without subtracting any blank value) to the exponential growth equation *Y(t)* = *b* + *Y*_0_^*^*e*^*kt*^ (using Prism 10 [GraphPad Software]), where *Y(t)* is the OD_600_ at time *t, b* is the OD_600_ of the well and empty medium (“blank”), *Y0* is the contribution of cells to OD_600_ at time *t* = 0, *k* is the growth rate (min-1), and *t* is the time (min).

### Whole Genome Sequencing

DNA was prepared using MasterPure (Epicentre) from 500 μL samples from overnight cultures of constructed clones or evolving populations grown at 30 °C or 37 °C in LB. Libraries for sequencing were prepared with Nextera DNA library and indexing kits (Illumina) and run in a MiSeq (Illumina). Sequencing reads were imported into CLC Genomics Workbench (Qiagen), and were trimmed to remove low quality reads and ambiguities, after which they were mapped to a reference genome containing all the modifications and known mutations. Indels and SNPs were detected using the low frequency variant detector, and structural rearrangements (duplications or deletions) were detected using visual scanning of mapped sequence read depth combined with the structural rearrangement tool in CLC.

## Acknowledgements

I thank Anna Knöppel for critical reading of the manuscript. This work was funded by the Swedish Research Council (VR-NT 2020-03512) and the Carl Trygger Foundation (CTS 22:2094).

## Notes

### Competing Interest Statement

The authors have declared no competing interest.

