## Supplemental text; Figures S1 - S4 for "Life with only 28 tRNAs: Reduced translation accuracy compensates for the lack of twelve tRNAs in *Salmonella enterica*"

Joakim Näsvall[1*]

[1] Dept. of Medical Biochemistry and Microbiology, Uppsala University, Uppsala, Sweden

This file contains:
Supplementary Text
Supplementary Figures S1 – S3

### Supplementary Text

#### Decoding specificities of tRNAs

For prediction of the codon reading patterns of tRNAs, the following wobble rules were applied (see Figure S1 for all S. enterica tRNAs):

1. tRNAs with cytidine (C) at the wobble position of the anticodon read only codons ending with guanidine (G)[1].
2. tRNAs with G at the wobble position read codons ending with C or uridine (U)[1].
3. tRNAs with 5-carboxymethoxyuridine (cmo$^5$U; uridine-5-oxyacetate) or related modified uridine derivatives at the wobble position read codons ending with adenine (A), G, U, and sometimes C[2–4].
4. tRNAs with 5-methylaminomethyluridine (mnm$^5$U) or related modified uridines read codons ending with A or G[5].
5. tRNAs with inosine (I, a modified A) at the wobble position read codons ending with U, C, and A[5].

Following these rules, fourteen *S. enterica* tRNAs were considered potentially nonessential as their cognate codons could be read by another tRNA (Table 1, Figure S1). However, among these we have previously identified one, $tRNA^{Thr}_{GGU}$, to be essential.

### Supplementary Figures

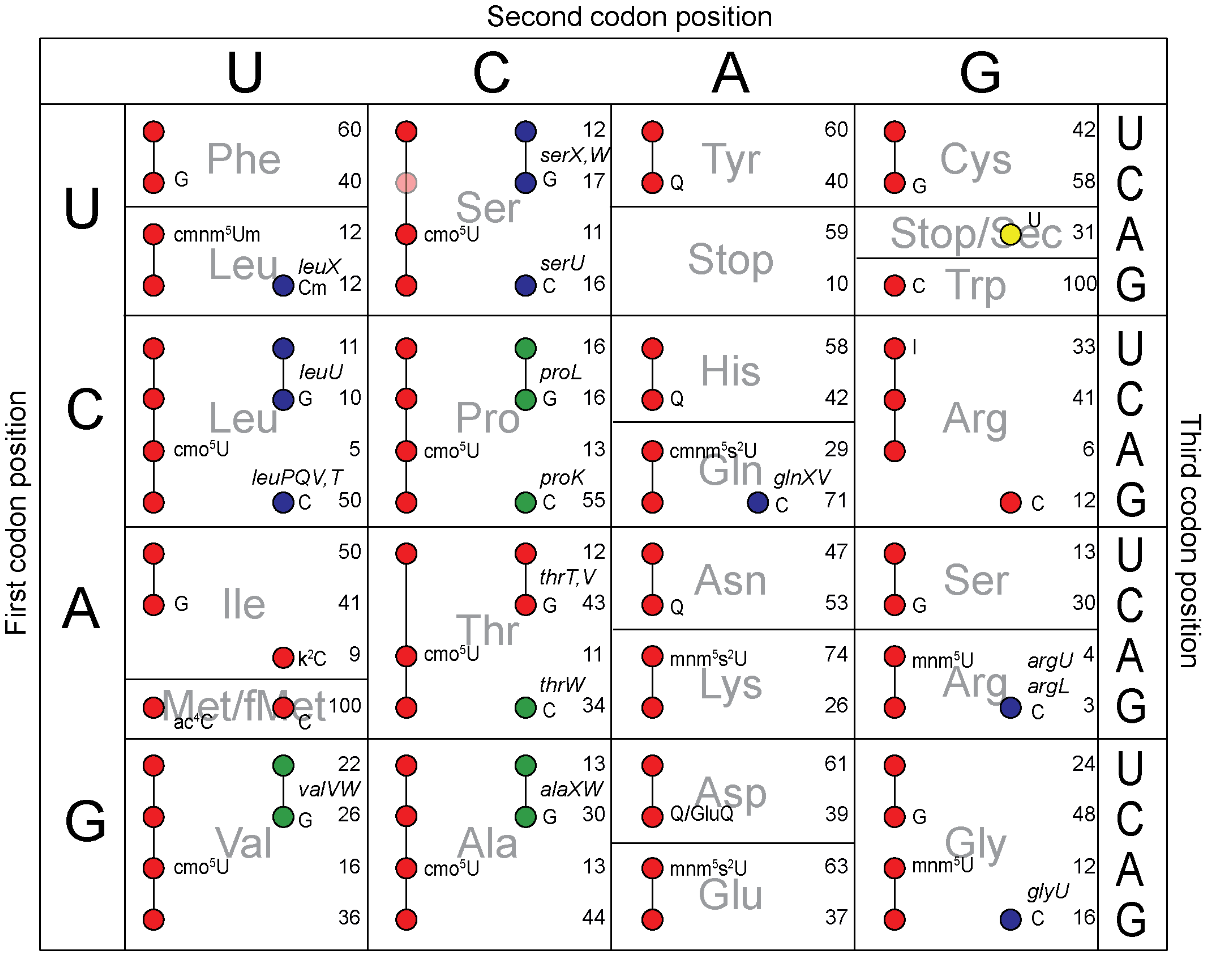


**Figure S1. Codon reading patterns of *S. enterica* tRNAs**. A colored circle indicate a tRNA capable of reading the corresponding codon. tRNAs capable of reading several codons are indicated by lines connecting several circles. For each tRNA, the nucleotide on the wobble position is indicated on the codon that is complementary to the anticodon of that tRNA; G, guanosine; Cm, 2´-O-methylcytidine; cmo$^5$U, 5-carboxymethoxyuridine; C, cytidine; k$^2$C, lysidine; ac$^4$C, N-4-acetylcytidine; Q, queuosine; GluQ, glutamylqueuosine; cmnm$^5$s$^2$U, 5-carboxymethylaminomethyl-2-thio-uridine; mnm$^5$s$^2$U, 5-methylaminomethyl-2-thio-uridine; U, uridine; I, inosine; mnm$^5$U, 5-methylaminomethyl-uridine. Red circles indicate tRNAs predicted to be essential based on being the only tRNA known to read one or more codons, green circles indicate nonessential tRNAs. The transparent red circle (UCC codon; tRNA$^{Ser}_{cmo5UGA}$) indicate it is unknown if this tRNA can read that codon. Blue circles indicate potentially nonessential tRNAs that read codons that could be read by another tRNA. The genes encoding the nonessential and potentially nonessential tRNAs are named next to those tRNAs.

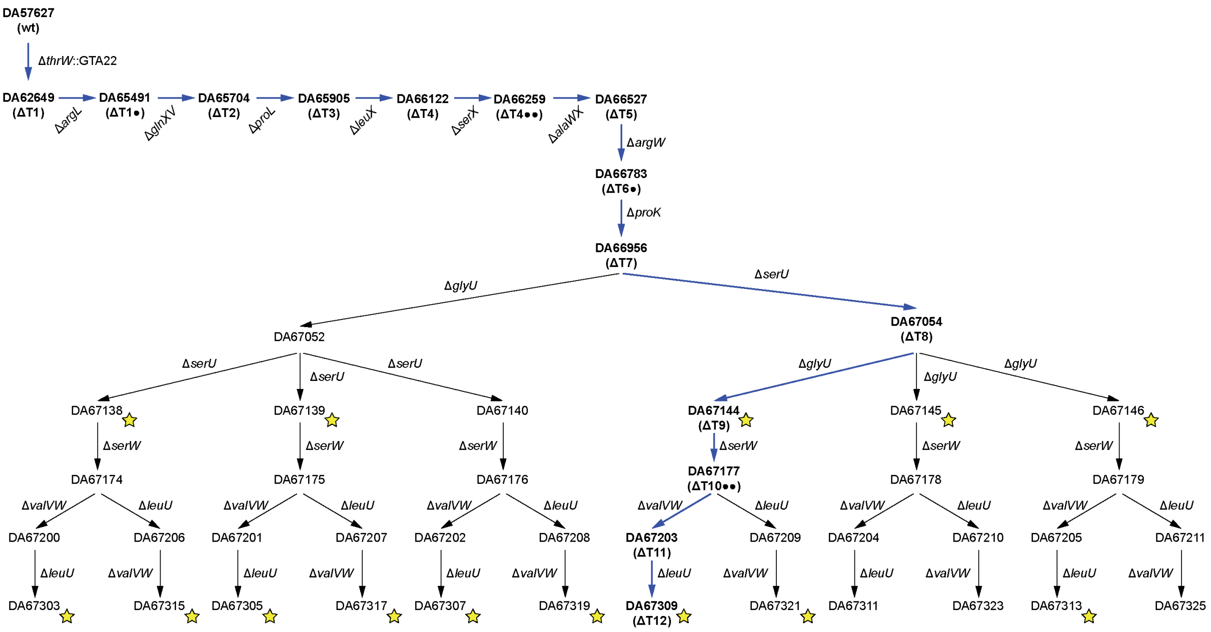


**Figure S2. Transduction series for generating a mutant lacking twelve nonessential tRNAs.** The deletions were introduced one after the other in the indicated order. The mutants in the direct ancestry of ΔT12 (blue arrows, bold text) are denoted by the most recently deleted tRNA gene followed by an indication of the number of missing tRNAs; *e.g.*, ΔT1 is missing one tRNA, ΔT2 is missing two, *etc.* •, *argL* and *argW* encode tRNAs with the same anticodons; ••, *serX* and *serW* encode identical tRNAs. Only completely missing tRNAs (all genes encoding tRNAs with the same anticodon deleted) add to the count of missing tRNAs. Stars indicate strains that were whole genome sequenced (Table S1).

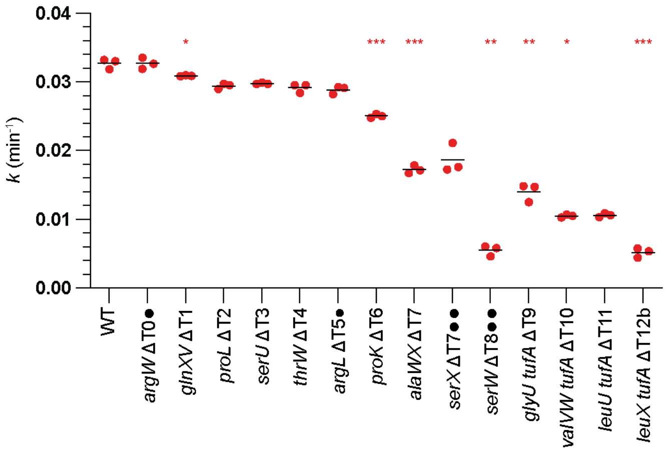


**Figure S3. Growth rates of tRNA deletion mutants at 37 °C.** The construction series leading up to ΔT12b; the mutants are denoted by the most recently deleted tRNA gene followed by an indication of the number of missing tRNAs; *e.g.*, ΔT1 is missing one tRNA, ΔT2 is missing two, *etc.* •, *argL* and *argW* encode tRNAs with the same anticodons; ••, *serX* and *serW* encode identical tRNAs. Only completely missing tRNAs (all genes encoding tRNAs with the same anticodon deleted) add to the count of missing tRNAs. Between transduction 10 (*serW*) and 11 (*glyU*) a mutation in *tufA* (EF-Tu) appeared, with a simultaneous growth rate improvement. Asterisks show significant differences according to a two-tailed Student's *t*-test where each mutant was compared to its parent (*i.e.*, ΔT12 was compared with ΔT11, which in turn was compared to ΔT10, *etc.*), *, $p < 0.05$; **, $p < 0.005$; ***, $p < 0.005$.

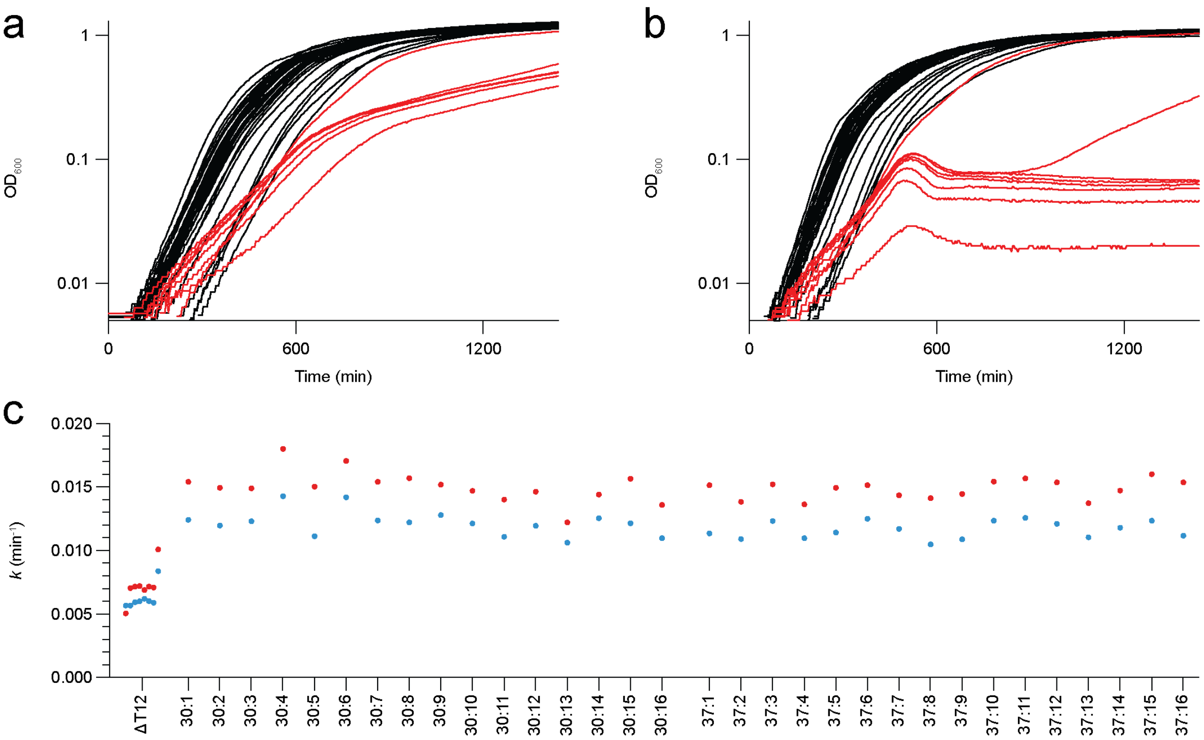


**Figure S4. Growth of evolved populations.** Batch growth curves at **(a)** 30 °C and **(b)** 37 °C. Black, evolved populations; red, ΔT12. **(c)** Exponential growth rates of evolved populations at 30 °C (blue) and 37 °C (red). The ancestor (ΔT12) was grown in overnight cultures at 30 °C as eight biological replicates and diluted 1:1000, while the populations were grown from 1:1000 dilutions of thawed freezer stocks. The same 1:1000 diluted cultures were split to two Bioscreen plates that were run at different temperatures. Note that the exponential phase ended at lower $OD_{600}$ for the ancestor than the the evolved populations, and six of the replicates of the ancestor stopped growing after approximately 500 minutes at 37 °C. Two of the replicates of the ancestor displayed accelerated growth (most obvious at 37 °C) after the end of the exponential phase, indicative of accumulation of pre-existing faster-growing mutants in those cultures.
